# Free energy landscape for the entire transport cycle of triose-phosphate/phosphate translocator

**DOI:** 10.1101/206912

**Authors:** Mizuki Takemoto, Yongchan Lee, Ryuichiro Ishitani, Osamu Nureki

## Abstract

Secondary active transporters translocate their substrates using the electrochemical potentials of other chemicals, undergoing large-scale conformational changes. Despite extensive structural studies, the atomic details of the transport mechanism still remain elusive. Here we performed a series of all-atom molecular dynamics simulations of the triose-phosphate/phosphate translocator (TPT), which exports organic phosphates in the chloroplast stroma in strict counter exchange with inorganic phosphate (P_i_). Biased sampling methods, including string method and umbrella sampling, successfully reproduced the conformational changes between the inward– and outward-facing states, along with the substrate binding. The free energy landscape of this entire TPT transition pathway demonstrated the alternating access and substrate translocation mechanisms, which revealed P_i_ is relayed by positively charged residues along the transition pathway. Furthermore, the conserved Glu207 functions as a “molecular switch”, linking the local substrate binding and the global conformational transition. Our results provide atomic-detailed insights into the energy coupling mechanism of antiporter.

## Introduction

Membrane transporters function as gatekeepers that permeate hydrophilic metabolites through a lipid bilayer. The secondary active transporters catalyze the uptake of compounds essential for cells or organelles or the elimination of unnecessary compounds, using the electrochemical potential of another compound. Generally, these active transporters achieve the transport of substrates by undergoing conformational transitions between the inward-facing (IF) state, the outward-facing (OF) state, and their intermediate, the occluded (Occ) state. This mechanism, called the alternating access mechanism(Drew & Boudker, 2016; Jardetzky, 1966), is universally required for all active transporters, since opening the gates on both sides of the membrane would permit the free diffusion of the substrate across the membrane and hinder the energy coupling. Although the energy coupling mechanism of membrane transporters is essential for all cells and organelles, the molecular basis of the energy coupling for almost all transporters remains elusive.

The plastidic phosphate translocator (pPT) family is one of the membrane transporter groups that exchanges phosphorylated C3, C5 and C6 carbon sugars with inorganic phosphate (P_i_) on the inner membrane of the plastid in plant and alga cells(Knappe, Flügge, & Fischer, 2003; Weber & Linka, 2011). The pPT family is further divided into four subfamilies, TPT, GPT, XPT and PPT, according to their substrate specificities for triose-phosphate, glucose-6-phosphate, xylulose-5-phosphate and phospho*enol*pyruvate, respectively(Eicks, Maurino, Knappe, Flügge, & Fischer, 2002; Fischer et al., 1997; Kammerer et al., 1998; Weber & Linka, 2011). The triose-phosphate/phosphate translocator (TPT) family in green plants exports triose phosphates and 3-phosphoglyceric acids (3-PGA) produced by photosynthesis and imports P_i_ into stroma U.-I. (Flügge et al., 1991; U. I. I. Flügge et al., 1989; Weber & Linka, 2011). Thus, TPT contributes to efficient carbon fixation and plant growth(Hattenbach, Muller-Rober, Nast, & Heineke, 1997; Häusler, Schlieben, Nicolay, et al., 2000; Häusler, Schlieben, & Flügge, 2000; Heineke et al., 1994; Riesmeier et al., 1993; Schneider et al., 2002). Each pPT subfamily catalyzes the strict 1:1 counter exchange of phosphorylated carbon compounds with P_i_ to ensure phosphate homeostasis between the stroma and the cytosol(Fliege, Flügge, Werdan, & Heldt, 1978; Weber, Schwacke, & Flügge, 2005). They can also mediate P_i_/P_i_ exchange in a reconstituted system, such as liposomes(Linka, Jamai, & Weber, 2008).

Recently, we reported the crystal structure of GsGPT from the thermophilic red alga *Galdieria sulphuraria*(Lee et al., 2017), which is functionally similar to TPT, rather than GPT, despite its name based on the sequence similarity(Linka et al., 2008). GsGPT is composed of 10 α-helical transmembrane (TM) helices (Figure 1a), and adopts the drug metabolite transporter (DMT) superfamily(Jack, Yang, & H. Saier, 2001) fold, similar to that of the recently reported bacterial metabolite transporter, *Starkeya novella* YddG (SnYddG)(Tsuchiya et al., 2016), in which the N-terminal half (TM1-5) is related to the C-terminal half (TM6-10) by two-fold pseudo-symmetry. We obtained the two crystal structures bound with different substrates, P_i_ and 3-PGA (Figure 1b and Figure 1—figure supplement 1). In both crystal structures, the phosphate moieties of the substrates are recognized by the well-conserved basic residues, Lys204, Lys362, and Arg363 (Figure 1b). This central substrate binding site is occluded from the solvent by hydrophobic residues, referred to as the inside and outside gates (Figure 1c). These observations indicated that the crystal structure represents a substrate-bound Occ state. The structural comparison with SnYddG in the OF state revealed the basis of the transport mechanism, in which the opening and closing of these gates regulate the IF and OF conformational changes(Lee et al., 2017). However, the detailed mechanisms of the conformational changes and the substrate binding/releasing by the DMT superfamily have remained elusive.

**Figure 1.**
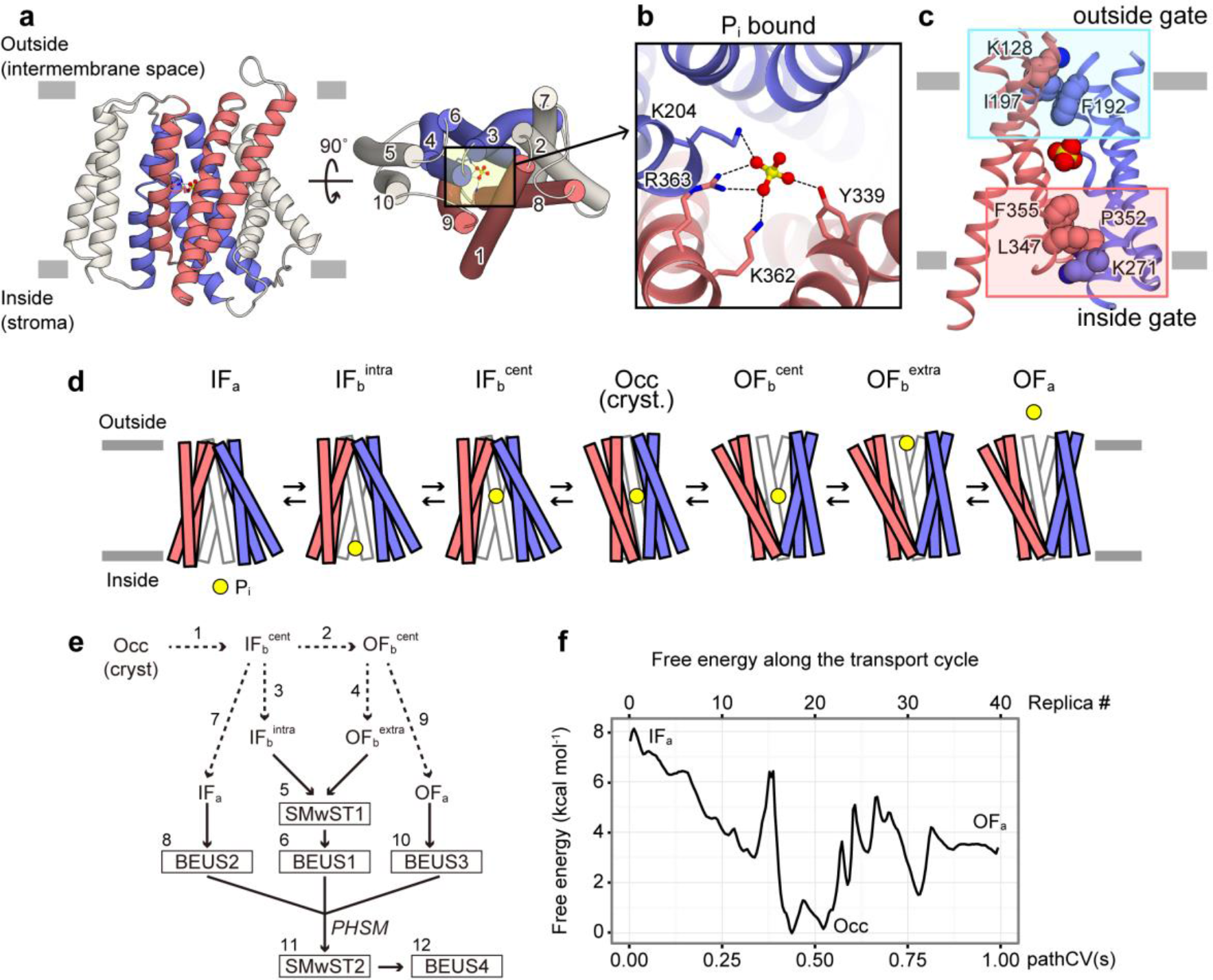
The crystal structure of GsGPT and simulation overview. (a) The crystal structure of GsGPT in the P_i_-bound state (PDB ID: 5Y78), viewed from the plane of the membrane (left) and from the intermembrane space (right). The numbers in the right panel represent the numbering of the TM helices. Hereafter, TM1, 8 and 9 are colored red, TM3, 4 and 6 are colored blue, and the others are colored white. (b) Close-up view of the central binding site in the P_i_-bound state. Key residues involved in substrate binding are shown in stick models. Dotted lines represent polar interactions. (c) Inside and outside gates of GsGPT. The inside and outside gates are highlighted within magenta and cyan rectangles, respectively. The substrate and gate-forming residues are shown in CPK models. TM2, 5, 7 and 10 are not shown. (d) Schematic representation of states, IF_a_ (IF-*apo*), IF_b_^intra^ (IF-bound at intracellular gate), IF_b_^cent^ (IF-bound at central binding site), Occ, OF_b_^cent^ (OF-bound at central binding site), OF_b_^extra^ (OF-bound at extracellular gate) and IF_a_ (OF-*apo*), involved in the currently studied transport cycle. P_i_ is represented by the yellow circle. The TM segments are colored in the same manner as in panel a. TM2 and 7 are not shown. (e) Graphical representation of the iterative scheme used for designing the simulations shown in Table 1. The number corresponds to the simulation numbering in Table 1. (f) Free energy profile along the simulated transport cycle shown in c, based on the final simulation (simulation set 12). pathCV(s) represents path collective variable s (equation (1))

**Table 1.** simulation set list.

| Sim # | Transition | Method | Collective variable | # of replicas x sim time |
| --- | --- | --- | --- | --- |
| 1 | $\text{cryst.} \rightarrow \text{IF}_b^{\text{cent}}$ | non-Eq. | $\Delta D$ | 10 ns |
| 2 | $\text{IF}_b^{\text{cent}} \rightarrow \text{OF}_b^{\text{cent}}$ | non-Eq. | $\Delta D$ | 20 ns |
| 3 | $\text{IF}_b^{\text{cent}} \rightarrow \text{IF}_b^{\text{intra}}$ | non-Eq | $Z_{\text{Pi}}$ | 10 ns |
| 4 | $\text{OF}_b^{\text{cent}} \rightarrow \text{OF}_b^{\text{extra}}$ | non-Eq | $Z_{\text{Pi}}$ | 10 ns |
| 5 | $\text{IF}_b^{\text{intra}} \rightleftharpoons \text{Occ} \rightleftharpoons \text{OF}_b^{\text{extra}}$ | SMwST | $\Delta D / Z_{\text{Pi}}$ | 40 x 1.2 ns = 48 ns |
| 6 | $\text{IF}_b^{\text{intra}} \rightleftharpoons \text{Occ} \rightleftharpoons \text{OF}_b^{\text{extra}}$ | BEUS | pathCV(s) | 20 x 20 ns = 400 ns |
| 7 | $\text{IF}_b^{\text{cent}} \rightarrow \text{IF}_a$ | non-Eq. | $Z_{\text{Pi}}$ | 25 ns |
| 8 | $\text{IF}_b^{\text{cent}} \rightarrow \text{IF}_a$ | BEUS | $Z_{\text{Pi}}$ | 16 x 20 ns = 320 ns |
| 9 | $\text{OF}_b^{\text{cent}} \rightarrow \text{OF}_a$ | non-Eq. | $Z_{\text{Pi}}$ | 25 ns |
| 10 | $\text{OF}_b^{\text{cent}} \rightarrow \text{OF}_a$ | BEUS | $Z_{\text{Pi}}$ | 16 x 20 ns = 320 ns |
| 11 | $\text{IF}_a \rightleftharpoons \text{Occ} \rightleftharpoons \text{OF}_a$ | SMwST | $\Delta D / Z_{\text{Pi}}$ | 40 x 5.5 ns = 220 ns |
| 12 | $\text{IF}_a \rightleftharpoons \text{Occ} \rightleftharpoons \text{OF}_a$ | BEUS | pathCV(s) | 40 x 50 ns = 2000 ns |
| | | | total | $\sim 3.4 \mu\text{s}$ |

Molecular dynamics (MD) simulations have been used to study the dynamics and conformational transitions of various membrane transporters(Khalili-Araghi et al., 2009; Shaikh et al., 2013). Our previous MD simulation, in which the substrate was removed from the crystal structure of GsGPT, showed the rapid conformational change from the Occ state to the IF and OF states, and provided insights into the transport mechanism of GsGPT(Lee et al., 2017). Generally speaking, a conventional MD simulation cannot sample a large and slow conformational change in the limited computational time, because of the high free energy barriers between the states. In fact, our previous MD simulation of the substrate-bound GsGPT did not reveal any significant conformational change in the 100-ns timescale(Lee et al., 2017). However, in the physiological transport cycle, spontaneous conformational changes between the Occ and IF/OF states, as well as the binding and release of the substrate (Figure 1d), actually occur in a much longer time scale, which is not accessible by the conventional MD simulation methods.

This problem can be solved by employing biased sampling methods, such as umbrella sampling(Torrie & Valleau, 1977). In this study, we used bias-exchange umbrella sampling (BEUS)(Moradi, Enkavi, & Tajkhorshid, 2015; Moradi & Tajkhorshid, 2014; Sugita, Kitao, & Okamoto, 2000), an umbrella sampling method combined with Hamiltonian replica exchange sampling(Sugita et al., 2000), which can perform more efficient sampling than normal umbrella sampling. To find the optimum transition pathway of the entire transport cycle of GsGPT, we employed the string method with swarms of trajectories (SMwST)(Pan, Sezer, & Roux, 2008) path-finding algorithm. By combining these sampling techniques, we achieved the *in silico* reconstruction of a feasible transport cycle for GsGPT in the explicit solvent and lipid bilayer, and we observed the substrate binding and releasing events for the first time for the DMT superfamily transporters. Comprehensive structural analyses revealed the strict 1:1 exchange mechanism of GsGPT in atomic detail. We found the unexpected common feature of the conformational transition mechanisms with the well-studied major facilitator superfamily (MFS) transporters(Quistgaard, Löw, Guettou, & Nordlund, 2016), despite their completely different protein folding. Along with the computational simulations, our functional analysis of the mutants provided experimental evidence supporting the MD simulation observations. Taken together, we revealed the atomistic molecular mechanism of GsGPT, which involves the structural coupling between the local conformational change induced by substrate binding and the global conformational transition of the transporter.

## Results

### Reconstruction of the GsGPT transport cycle

To reconstruct the GsGPT transport cycle, we first tried to find the reaction coordinates (collective variables) to induce the conformational transition of GsGPT. Various trials revealed that two distance parameters, D_in_ and D_out_, were sufficient to induce the transition to the IF and OF states from the Occ state crystal structure (Figure 1—figure supplement 2b-e), where D_in_ and D_out_ represent the center of mass distances between the Cα atoms of the inside and outside gate residues, respectively (see Methods for a detailed definition). Moreover, the one-dimensional collective variable, ΔD = D_out_ − D_in_, can also be used to induce the conformational change from IF to OF (Figure 1—figure supplement 2f). For simplicity, we used this ΔD to describe the conformational space of GsGPT. Another reaction coordinate for the transition cycle is the position of P_i_ along the membrane normal axis, referred to as Z_Pi_ (Figure 1—figure supplement 2a). Finally, we performed 12 sets of biased samplings in the two-dimensional collective variable space, ΔD and Z_Pi_.

The reconstruction of the transport cycle is performed by the following steps (Figure 1e): first of all, we performed the non-equilibrium pulling to induce the transitions from Occ to IF_b_^cent^ and OF_b_^cent^ (IF– and OF-bound at the central binding site, respectively) (Figure 1d), using the collective variable ΔD (simulation sets 1 and 2 in Table 1). The subsequent non-equilibrium pulling for Z_Pi_ was performed to generate the structures in which P_i_ is bound near the gate, IF_b_^intra^ (IF-bound at intracellular gate) and OF_b_^extra^ (OF-bound at extracellular gate) (simulation sets 3 and 4). To search the minimum free energy pathway of the IF_b_^intra^ ↔ Occ ↔ OF_b_^extra^ transition in the ΔD - Z_Pi_ two-dimensional space, we employed the string method with swarms of trajectories (SMwST)(Pan et al., 2008) (simulation set 5), followed by the free energy calculation with a set of bias exchange umbrella sampling (BEUS)(Moradi et al., 2015; Moradi & Tajkhorshid, 2014; Sugita et al., 2000) along the refined transition string (simulation set 6) using “path collective variables” (pathCV(s) and pathCV(ζ); equations (1) and (2))(Branduardi, Gervasio, & Parrinello, 2007). Next, to obtain the *apo* states of GsGPT, IF_a_ and OF_a_ in Figure 1c, we pulled the P_i_ from the IF_b_^cent^ and OF_b_^cent^ states to the bulk solvent region (simulation sets 7 and 9), and performed free energy calculations along the Z_Pi_ variable (simulation sets 8 and 10). The results of these three sets of free energy calculations were combined with the recently developed *post-hoc* string method (PHSM)(Moradi et al., 2015) to estimate the minimum free energy pathway for the full transition cycle, IF_a_ ↔ Occ ↔ OF_a_. The extracted transition pathway was further refined with SMwST (simulation set 11). Finally, the free energy calculation for the entire transport cycle (simulation set 12) was performed along this optimized transition pathway, using pathCV(s) and pathCV(ζ). We obtained the entire transport cycle of GsGPT with 1,000,000 configurations, and calculated the free energy along the optimum transition pathway using the weighted histogram analysis method (WHAM)(Kumar, Rosenberg, Bouzida, Swendsen, & Kollman, 1992) (Figure 1f and Video 1). (For detailed parameters of the simulations, see Methods.)

### Overall conformational change of GsGPT

To investigate the overall conformational change of GsGPT, representative structures corresponding to the free energy basins of the IF_a_ and OF_a_ states (Figure 1f) were compared with that of the Occ state (Figure 2a, b). The major differences between the IF and OF states were the outward-gate distance between the hairpin of TM3-4 and TM1 (Figure 2a), and the inward-gate distance between the hairpin of TM8-9 and TM6 (Figure 2b). The superimposition of the IF_a_ and OF_a_ states onto the Occ state revealed that the TM bundles composed of TM1, 8 and 9 (bundle1) and TM6, 3 and 4 (bundle2) are directly involved in the overall conformational change. In contrast, the other helices (TM2, 5, 7 and 10) do not participate in the open/closed conformational changes, and function as a scaffold for the conformational changes of bundle1 and bundle2. Hereafter, we refer to bundle1 and bundle2 (*i.e.*, TM1, 3, 4, 6, 8, and 9) as the core domain.

**Figure 2.**
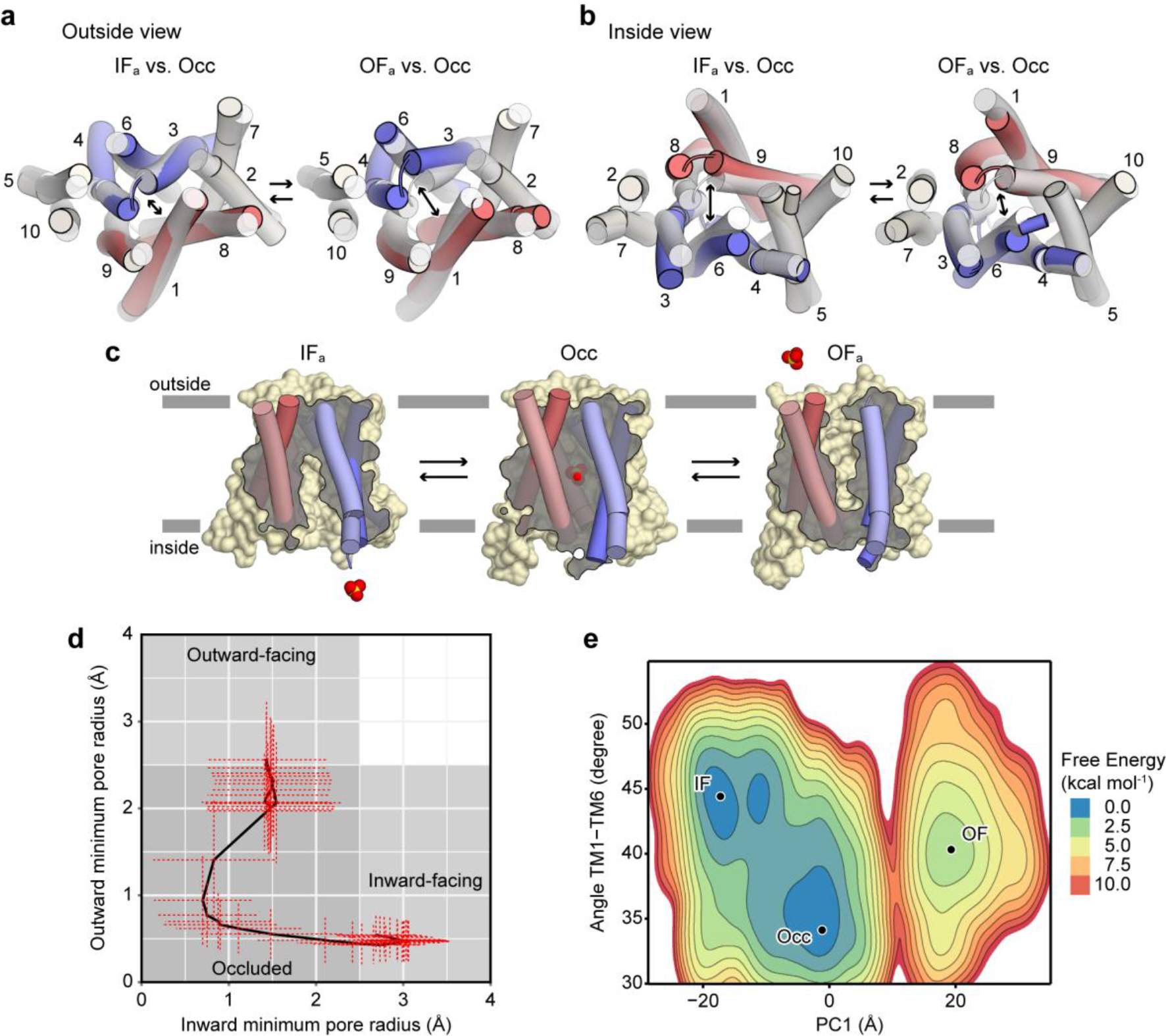
Overall conformational change. (a) Outside and (b) inside views of representative structures in the OF and IF states, shown in cylinder representations. Each structure was superimposed onto the Occ state structure, shown with a transparent cylinder. Bundle1 and bundle2 are colored red and blue, respectively, and the other helices are colored white. The number represents the numbering of the TM helices. (c) Cross-sections of surface representations for the representative structures from the IF, Occ and OF states. P_i_ is shown in a CPK model. (d) Pore radii at the outside and inside gates. The mean values of the outside and inside radii in each replica were plotted. The error bars represent the standard deviation in each replica. (e) Free energy landscape in terms of (PC1, TM1-TM6 angle) space. PC1 represents the first principal component of all Cα atoms, and the inter-helical angle between TM1 and TM6 was calculated using the roll axis of the helix (obtained from the principal axis component analysis for all Cα atoms).

The molecular surfaces for the representative structures illustrate that the substrate binding site is exposed to opposite sides of the membrane in an alternating fashion, known as the alternating access mechanism(Drew & Boudker, 2016; Jardetzky, 1966) (Figure 2c). To confirm whether all of the structures obtained from the simulation satisfy the alternating access mechanism, we performed pore radius calculations along the transport pathway by HOLE(Smart, Neduvelil, Wang, Wallace, & Sansom, 1996) for all structures obtained from the simulation (for representative results, see Figure2—figure supplement 1). The outward and inward minimum radii were defined as the minimum radii of the pore in the regions of −20<z<−7 and 7<z<20, respectively, where z represents the coordinate along the pore pathway (z = 0 corresponds to the central binding site, and z > 0 corresponds to the outward direction). The plot of the outward *vs*. inward minimum radii illustrates that the conformational transition between the IF and OF states always visits the Occ-state region, and none of the structures visit the region where the substrate binding site is exposed to both the inward– and outward-side solvent (Figure 2d). This result indicates that the obtained transition pathway satisfies the alternating access, suggesting the validity of the simulated transition pathway.

Next, to reveal the mechanism of the overall conformational change, we investigated the internal conformational change within the core domain helices, bundle1 and bundle2. Bundle1 and bundle2 in the IF_a_ conformation both superimpose well onto those in the OF_a_ conformation, respectively (Figure 2—figure supplement 2a, b), suggesting the semi-rigid body movement of the core domains along the transport cycle. In fact, the distributions of the root-mean-square deviation (RMSD) values of the bundle1 and bundle2 Cα atoms against the crystal structure illustrated that the RMSD values for only bundle1 or bundle2 remained lower than that of bundle1+2 over the entire transport cycle (Figure 2—figure supplement 2c). These results suggest that the conformational transition between IF and OF is governed by the semi-rigid body tilting motions of bundle1 and bundle2, rather than the internal bending and straightening motion of the TM helices. To confirm this mechanism, we projected the free energy onto the two-dimensional space consisting of the first principal component of the Cα atoms, or PC1 (Figure 2—figure supplement 3a-c), and the inter-helical angle between TM1 and TM6 (Figure 2e). The PC1 correlates well with the ΔD, pathCV(s), and replica number (Figure 2—figure supplement 3d-f), and represents the global conformational transition as well as the ΔD. The inter-helical angle between TM1 and TM6 represents the inter-domain angle between bundle1 and bundle2. This free energy landscape shows three distinct free energy basins corresponding to the IF, Occ and OF states, suggesting that the tilting motions of bundle1 and bundle2 play a major role in the overall conformational change.

These results are in contrast with the previously suggested mechanism hypothesized from the static crystal structures of SnYddG and GsGPT, in which the bending motion of the TM helices governs the conformational transition(Lee et al., 2017). In the previous report of the bacterial DMT superfamily transporter, SnYddG, the bending and straightening of the core bundle, especially TM3-4 and TM8-9, were suggested to regulate the opening and closing of the inside and outside gates(Tsuchiya et al., 2016). However, the present results suggest that the tilting motion of the semi-rigid core domain, which resembles the rocker-switch mechanism proposed for the MFS transporters(Huang, Lemieux, Song, Auer, & Wang, 2003; Law, Maloney, & Wang, 2008), is sufficient to explain the conformational transition of GsGPT. Thus, the alternating access mechanism of the DMT superfamily is unexpectedly similar to that of the MFS transporters, despite their completely different TM topologies.

### Hydrogen bonds at the inside and outside gates are dispensable for the conformational change

In the crystal structure of GsGPT, the well conserved Lys271 and Lys128 residues form hydrogen bonds with the main chain carbonyl atoms on the opposite bundle, and lock the inside and outside gates, respectively (Figure 3a). In our previous analysis based on the structural comparison with SnYddG, the disruption and formation of these hydrogen bonds were suggested to regulate the gate opening and closing(Lee et al., 2017). However, in the present simulation, the two hydrogen bonds of Lys128 shown in Figure 3a were not always maintained, even in the IF state (Figure 3b; cyan), and the two hydrogen bonds of Lys271 were also not always maintained, even in the OF state (Figure 3b; magenta). Furthermore, the liposome-based transport assays of K271A, K128A and their double mutant revealed that all of these mutants still retain the P_i_/ P_i_ homo-exchange activity, as compared with the previously reported mutants in the substrate binding site, K204A, K362A and R363A(Lee et al., 2017) (Figure 3c). These results suggest that the hydrogen bond formation by Lys271 or Lys128 is not crucial for the conformational change. Our investigations are consistent with the previous functional analysis of the GsGPT homologue, the apicoplast phosphate translocator of *Toxoplasma gondii* (TgAPT), which revealed that the K67A mutation of TgAPT, corresponding to the K128A mutation of GsGPT, retained 10-20% activity(Brooks et al., 2010). Our simulation suggests that Lys128 and Lys271 play another role in the conformational transition of GsGPT, rather than the regulation of gate opening and closing by hydrogen bond formation.

**Figure 3.**
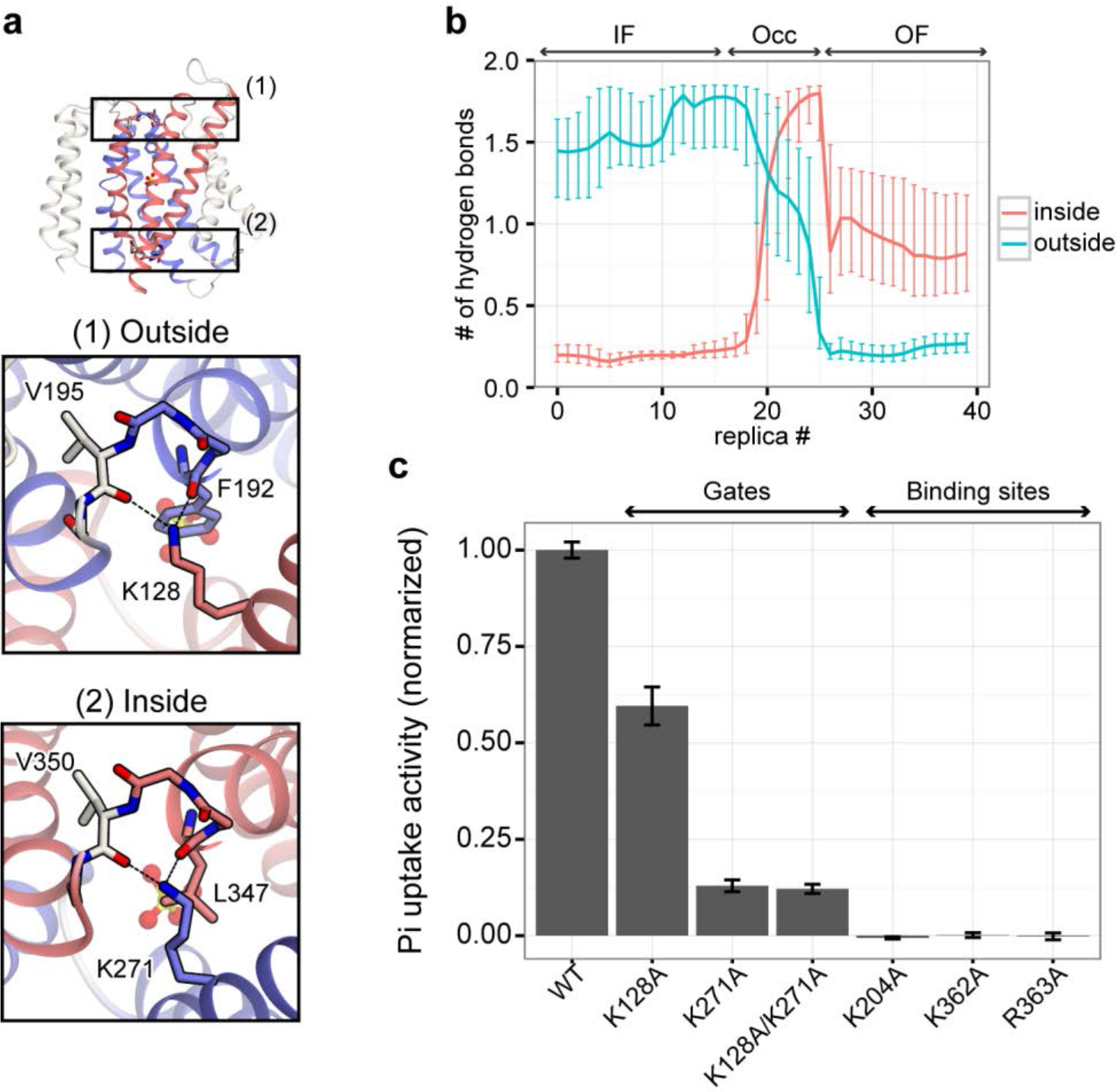
Hydrogen bonds at the inside and outside gates. (a) Overall structure of GsGPT (upper) and close-up views of the outside (center) and inside (lower) gates. The dotted line represents a hydrogen bond between a Lys residue and a main chain carbonyl atom. (b) The number of hydrogen bonds in the inside and outside gates (magenta and cyan, respectively), shown in panel a. The median value in each replica is plotted. Error bars represent the interquartile range (IQR). (c) Liposome-based mutational analysis. The levels of [^32^P]-P_i_ uptake by GsGPT mutants were normalized with the mean value of the wild-type. The error bars represent s.e.m. (n=3-9; see Figure 3—Source Data1). The values of K204A, K362A and R363A are from our previous study(Lee et al., 2017).

### Substrate binding and translocation mechanism

To gain detailed insight into the mechanism for substrate binding and translocation, we projected the free energy onto the PC1– Z_Pi_ plane, and compared the representative structures of the free energy basins (Figure 4). The translocation of P_i_ from the outside (intermembrane side) to the inside (stroma side) can be described by the following reaction steps (Video 1). (i) At first, P_i_ in the outside solution is captured by the positively charged residues, and passed to Lys128 (Figure 4a and Figure 4—figure supplement 1a). (ii) In addition to Lys128, P_i_ is captured by Lys204, one of the central substrate binding residues (Figure 4b and Figure 4—figure supplement 1b). (iii) The salt bridge between P_i_ and Lys128 is disrupted, and then P_i_ binds to Lys362 and Arg363 (Figure 4c and Figure 4—figure supplement 1c, d). (iv) The hydrogen bond between P_i_ and Tyr339 on TM9 is formed (Figure 4d and Figure 4—figure supplement 1e), which enables the outward halves of the core domains to approach each other, and the conformational change from the OF to Occ states is facilitated. (Figure 4d). This configuration is the most stable state in our simulation (Figure 1f and Figure 4), consistent with the fact that the two crystal structures of GsGPT were obtained in the substrate-bound Occ state. Note that two distinct configurations were degenerate in this Occ state in terms of the binding modes of Arg266: in one state, Arg266 binds to P_i_ with a water-mediated indirect interaction, as observed in the crystal structure (Figure4—figure supplement 2a); in the other, Arg266 directly forms a salt bridge with P_i_ (Figure 4—figure supplement 2b). These two states can be discriminated by the slight difference of Z_Pi_, and this difference might determine the direction of the conformational transition from the Occ state to the IF or OF state. (v) The inside gate is disrupted, and the conformational change from the Occ to IF state occurs (Figure 4e). (vi) P_i_ is dissociated from the central binding site and bound to both Arg266 and Lys271 (Figure 4f and Figure 4—figure supplement 1f, g). (vii) The salt bridge between P_i_ and Arg266 is disrupted, and P_i_ is released to the inside solution mediated by Lys271 and other basic residues located on the inward surface, *e*.*g*. Lys400 (Figure 4g). Each free energy basin corresponds to a distinct substrate binding mode, and the transport mechanism can be explained as the relaying of P_i_ by these pore lining, basic residues (except for Tyr339). The free energy barriers between the P_i_ binding modes correspond to the energy required for disrupting the salt bridge between P_i_ and these basic residues.

**Figure 4.**
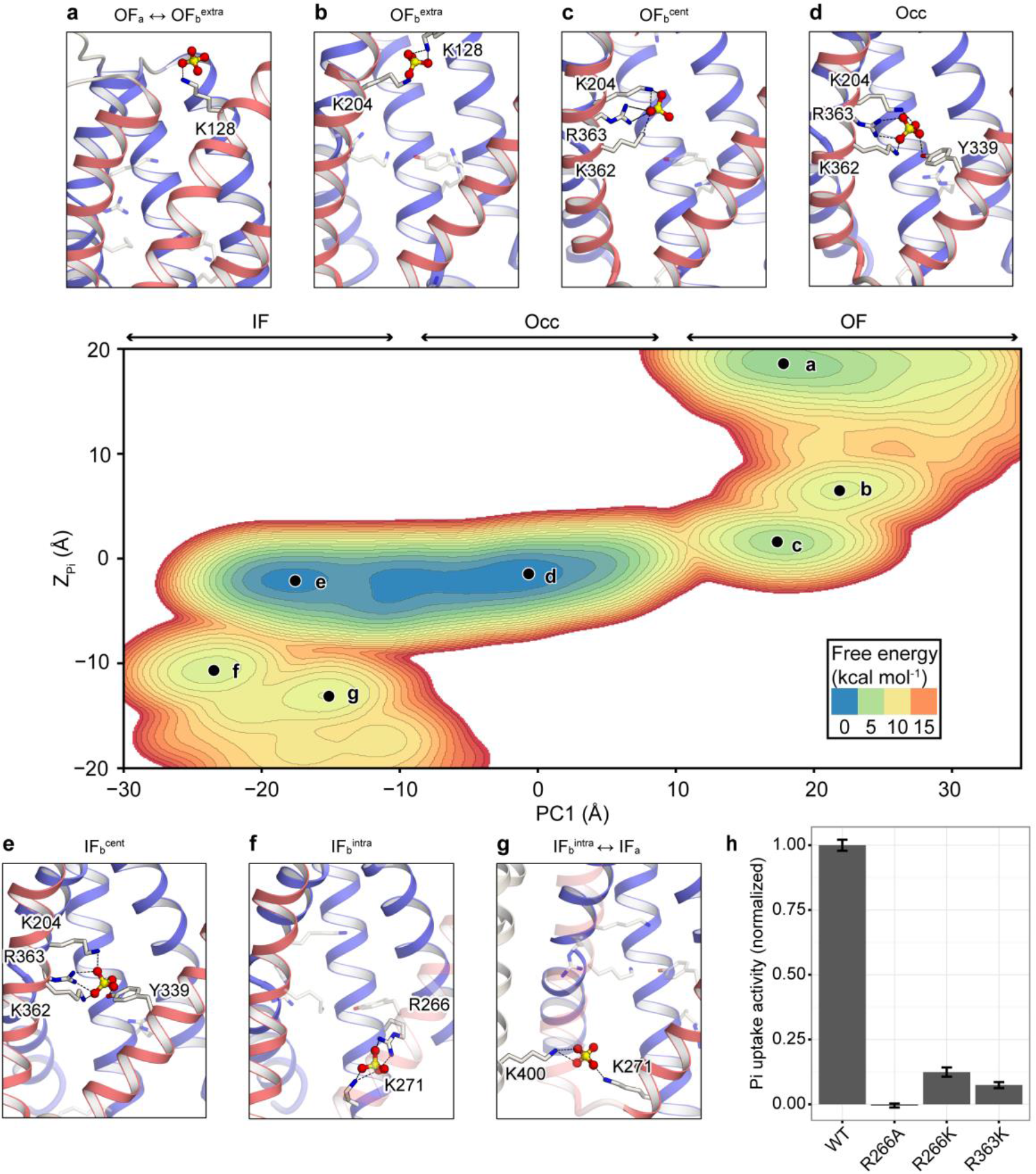
Substrate binding and global conformational change coupling. Free energy landscape in the (PC1, Z_Pi_) space is shown for the regions in which P_i_ is bound to GsGPT. (a-g) Snapshots of the binding site conformation corresponding to each free energy basin. The residues that directly interact with P_i_ in the transport cycle are shown with stick models. Dotted lines represent strong interactions in which the distance between P_i_ and each residue is less than 3.0 Å. TM1 is not shown, except in panel a. (h) Liposome-based mutational analysis. The levels of [^32^P]– P_i_ uptake by GsGPT mutants were normalized with the mean value of the wild-type. The error bars represent s.e.m. (n=3-9; see Figure 3—Source Data1).

The minimum distance plot between P_i_ and these pore lining residues shows that the residues in the central binding site, Lys204, Tyr339, Lys362 and Arg363, interact strongly with the substrate in all of the OF, Occ and IF states (Figure 4—figure supplement 1b-e). This observation is consistent with our previous mutational analysis, in which the mutations of these residues, K204A, Y339F, K362A and R363A, abolished the P_i_/P_i_ homo-exchange activity(Lee et al., 2017). It is also interesting to note that Lys271 and Lys128 in the inside and outside gates strongly interact with P_i_ in the transport cycle. Given that the hydrogen bond formation of Lys128 and Lys271 is dispensable for the conformational transition (Figure 3), the decreases of the transport activity in the K128A, K271A and K128A/K271A mutants may be due to the loss of these interactions. Thus, we can conclude that Lys128 and Lys271 primarily function as the initial binding sites for the substrates, rather than in the inside and outside gate formation. To further support this relaying mechanism of P_i_, we performed an additional mutational analysis of GsGPT using the liposome-based assay (Figure 4h). The result revealed that the R266A mutant also completely abolishes the P_i_ exchanging activity, showing the importance of Arg266 for the transport mechanism. In addition, the conservative mutants, R266K and R363K, exhibited significantly decreased activities (Figure 4h), suggesting that the shapes of these side chains are also important for the recognition of the tetrahedral arrangement of the substrate.

### Coupling mechanism of substrate binding and protein conformational change

To reveal the conformational coupling mechanism of GsGPT, we focused on Glu207 on TM4, which is the only conserved acidic residue in the positively-charged substrate binding pocket (Figure 5a). Calculations of the minimum distances between the neighboring basic residues, Lys204 and Arg363, revealed that Glu207 exchanges its salt bridge partner during the transport cycle (Figure 5a, b). In the IF_a_ state, Glu207 mainly forms an intra-helical salt bridge with Lys204 (Fig 5b; magenta line). This intra-helical salt bridge fixes the side chain of Lys204 to the Glu207 side, thereby causing the electrostatic repulsion between Lys204 and Arg363 to prevent the conformational transition to Occ and stabilize the IF conformation (Figure 5a; left). By contrast, in the OF_a_ state, Glu207 forms a stable inter-helical salt bridge with Arg363 on TM9 (Fig 5b; cyan line). This inter-helical salt bridge between Glu207 and Arg363 clamps the inward halves of TM4 and TM9 together (Figure 5a; right), thereby fixing the arrangement of bundle1 and bundle2 to the OF conformation. P_i_ binding to these basic residues weakens the salt bridges between the basic residues and Glu207, which may facilitate the exchange of the salt bridge partner of Glu207 (Figure 5a; center).

**Figure 5.**
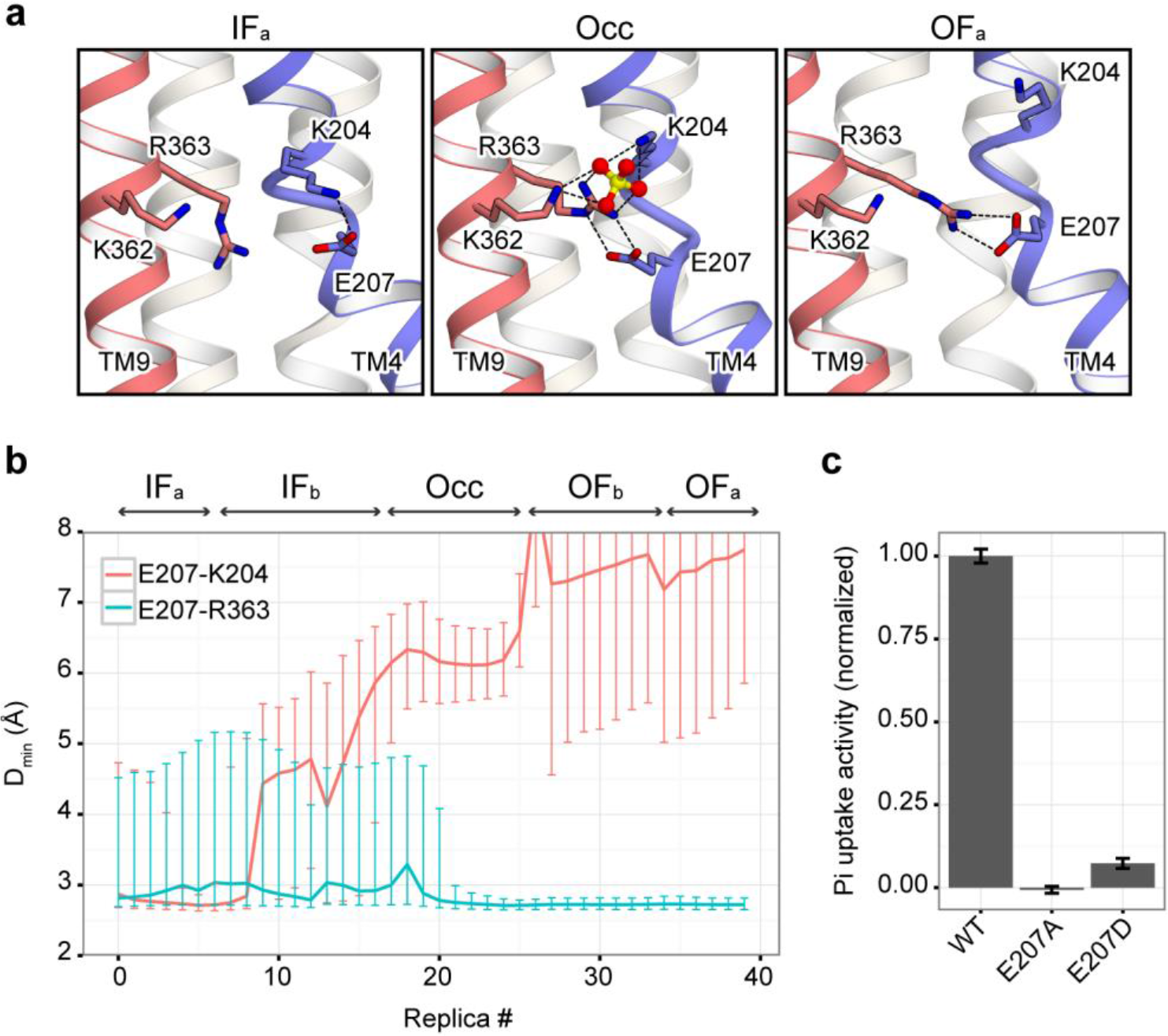
E207 in the substrate binding site. (a) Close-up view of the substrate binding site for representative structures of IF, Occ and OF. Dotted lines represent strong salt bridge interactions, in which the distance between each atom is less than 3.5 Å. (b) Minimum distance between Glu207 and Lys204 (magenta) and between Glu207 and Arg363 (cyan). The median value of the distance in each replica is plotted, and the error bars represent the interquartile range (IQR). (c) Liposome-based mutational analysis. The levels of [^32^P]-P_i_ uptake by GsGPT mutants were normalized with the mean value of the wild-type. The error bars represent s.e.m. (n=3-9; see Figure 3—Source Data1).

This salt-bridge rearrangement involving Glu207 leads to a plausible mechanism of the conformational coupling, which can explain how the local conformational change upon the substrate binding is amplified to the global motions of bundle1 and bundle2. In this mechanism, Glu207 plays a role as a “switch” that turns on the global tilting motions of TM4 and TM9 upon substrate binding. Since TM4 and TM9 constitute the (semi-)rigid bodies of bundle1 and bundle2, respectively, these tilting motions of TM4 and TM9 directly result in the global IF/OF conformational change of GsGPT (Figure 5—figure supplement 1). This mechanism is in contrast to our previous working model based on the static crystal structure, in which the electrostatic repulsion by the positively charged residues in the substrate binding pocket is simply neutralized by the binding of the negatively-charged substrate, thereby enabling the global IF/OF conformational change(Lee et al., 2017). To further support the importance of Glu207, we performed liposome-based assays of Glu207 mutants. The results revealed that the E207A mutation abolishes the P_i_/P_i_ homo-exchange activity (Figure 5c), in spite of its stable membrane expression (Figure 3—figure supplement 2). Moreover, the conservative mutation, E207D, significantly reduces the exchange activity (Figure 5c), probably due to the insufficient side chain length to form the salt bridge network (Figure 5a). Taken together, in addition to the previous mechanism based on the electrostatic repulsion and neutralization, the switch mechanism by Glu207 plays a pivotal role in the coupling mechanism of the local and global conformational changes, thus allowing the strict 1:1 exchange of the substrates by GsGPT.

## Discussion

Our simulation, combining string method and umbrella sampling, successfully reconstructed the entire transport cycle of GsGPT from a single crystal structure. The computational analysis for all configurations obtained from the simulation revealed the antiport mechanism of GsGPT in atomic detail.

In the current study, we used a highly symmetric substrate, P_i_, to simplify the collective variable used in the simulations. However, we can gain insight into other pPT family protein substrates that have more complicated structures, such as 3-PGA. Although pPT family proteins transport various types of phosphosugars, depending on their subtypes(Lee et al., 2017; Weber & Linka, 2011), all of the substrates have a common structural feature: one oxygen atom of P_i_ is replaced with a sugar moiety. In our simulation, the substrate binding pocket in all of the binding modes can accommodate a compound in which one oxygen atom of P_i_ is replaced with a sugar moiety (Figure 4a-g), as observed in our previous 3-PGA bound crystal structure(Lee et al., 2017). In addition, our previous study revealed that the recognition and discrimination of the sugar moiety of the substrates is achieved by the side chains located on the opposite side of the P_i_ binding residues, such as His185 (Figure 1—figure supplement 1b), which are not involved in the currently proposed transport mechanisms. The conformational transition is regulated by the binding of the phosphate moiety, and the binding manner of the phosphate moiety is the same in both the P_i_ and 3-PGA bound crystal structures(Lee et al., 2017) (Figure 1 and Figure 1—figure supplement 1). Therefore, it is highly likely that the transport mechanism of other substrates by GsGPT is similar to that of P_i_ observed in our simulations. Moreover, given that the P_i_ binding residues observed in our simulation (Lys128, Lys204, Arg266, Lys271, Tyr339, Lys362 and Arg363) are well conserved among the pPT family members(Lee et al., 2017), and that the free energy landscape is governed almost entirely by the binding mode of P_i_ to these conserved residues, all of the transporters belonging to the pPT family may have similar free energy landscapes to that of GsGPT (Figure 4). Thus, our results can be generalized to the transport mechanism for all pPT family transporters.

Our previous research on the bacterial DMT superfamily protein, SnYddG, revealed that the best-characterized small drug resistance (SMR) family protein, *E. coli* EmrE(Chen et al., 2007; Henzler-Wildman, 2012), shares a similar TM topology with SnYddG, suggesting an evolutional relationship between SMR and other DMT proteins(Tsuchiya et al., 2016). EmrE forms a dimer of four TM segments, and exchanges protons and cationic drugs(Chen et al., 2007; Henzler-Wildman, 2012). The structural comparison of EmrE with SnYddG and GsGPT revealed that EmrE lacks the TM helices corresponding to TM2 and TM7 in SnYddG and GsGPT(Tsuchiya et al., 2016). Our simulation showed that these TM helices work as a scaffold, and are not directly involved in the conformational transition. Thus, the transport mechanism of EmrE can also be explained as the rocker-switch movement of a pair of three TM bundles (TM1, 2 and 3 of each protomer), corresponding to bundle1 and bundle2 of GsGPT (Figure 2c).

It is worth comparing the conformational regulation mechanism of GsGPT with that of GlpT, which belongs to MFS and functions as a glycerol-3-phosphate/P_i_ exchanger(Huang et al., 2003; Law, Maloney, et al., 2008). GlpT has a different evolutionary origin and protein folding from those of GsGPT, but is functionally similar to GsGPT. The previous biochemical and MD simulation studies(Law, Almqvist, et al., 2008; Moradi et al., 2015) revealed that the inter- and intra-domain salt bridges between two basic (R45 and K46) and two acidic (D274 and E299) residues stabilize the different conformations during the transport cycle, and substrate binding weakens these interactions, allowing the conformational transition. In the case of GsGPT, our simulations revealed that the inter- and intra-helical salt bridges involving Glu207, which connect bundle1 and bundle2, stabilize the IF and OF conformations (Figure 5), and these interactions are weakened by substrate binding. Furthermore, the overall conformational change is described by the rocker-switch motions of bundle1 and bundle2 (Figure 2), which are widely observed in MFS transporters, including GlpT(Huang et al., 2003; Law, Maloney, et al., 2008). Therefore, the conformational change mechanism by the rocker switch motion and the conformational regulation mechanism by the salt bridge formation and disruption in GlpT and GsGPT are quite similar to each other. These similarities between GlpT and GsGPT over protein superfamily, in terms of both functions and mechanisms, may represent an example of the convergent evolution of the transporter mechanisms.

## Methods

### Simulation system setup

All MD simulations were performed with NAMD 2.9-2.11(Phillips et al., 2005), along with its Colvars Module(Fiorin, Klein, & Hénin, 2013). The simulation system was 96×96×100 Å^3^, and contained GsGPT (PDB ID: 5Y78), a molecule of P_i_, a POPC bilayer, 150 mM NaCl, and TIP3 water molecules. All of the water molecules that were visible in the crystal structure were kept. The missing hydrogen atoms were added by the psfgen plugin of VMD(Humphrey, Dalke, & Schulten, 1996). The net charge of the system was neutralized by adding 150 mM NaCl. The molecular topology and force field parameters from Charmm36 were used(Best et al., 2012; Klauda et al., 2010). The systems were first energy minimized for 1,000 steps with fixed positions of the non-hydrogen atoms, and then for another 1,000 steps with 10 kcal/mol restraints for the non-hydrogen atoms, except for the lipid molecules within 5.0 Å from the protein. Next, equilibrations were performed for 0.1 ns under NVT conditions, with 10 kcal mol^−1^ Å^−2^ restraints for the heavy atoms of the protein. Finally, equilibration was performed for 1.0 ns under NPT conditions with the 1.0 kcal mol^−1^ Å^−2^ restraints for all Cα atoms of the protein, followed by a 100 ns equilibration without the restraints. The equilibrated system was further used for the next simulation, as indicated in Table 1 and below. For all simulations in Table 1, the SHAKE algorithm was used to constrain all hydrogen-containing bonds to enable a 2 fs timestep. Constant temperature was maintained with Langevin dynamics at 310 K, and constant pressure was maintained with a Nosé -Hoover Langevin piston(Feller, Zhang, Pastor, & Brooks, 1995) at 1 bar. Electrostatic interactions were calculated by the particle mesh Ewald method(Darden, York, & Pedersen, 1993).

### Collective variables

In order to induce the conformational change from the Occ state crystal structure, we used the collective variables of ΔD and Z_Pi_. ΔD was defined as follows: ΔD = D_out_ − D_in_, where D_out_ was defined as the distance between the Cα mass centers of the two residue groups, 126-130 and 191-200, and D_in_ was defined as the distance between the Cα mass centers of the two residue groups, 269-273 and 347-355. Z_Pi_ was used to track the position of the substrate P_i_, and defined as the projection of the distance vector between the phosphorus atom of P_i_ and the reference point onto the membrane normal axis. The reference point of Z_Pi_ was defined as the center of mass of the Cα atoms of the P_i_ binding residues in the crystal structure, *i*.*e*., K204, K362 and R363. In all of our simulations, P_i_ was also restrained within a cylinder with a 10 Å radius, which was aligned to the membrane normal axis and centered to the mass center of the protein, using a half-harmonic potential with a spring constant of 10 kcal mol^−1^ Å^−2^.

To sample the configurations for the entire transport cycle of GsGPT, we used the following strategy. Generally, it is difficult to sample sufficiently in an *n*-dimensional collective variable space, 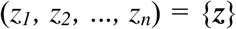, due to the computational cost. A practical solution is to perform the sampling only around a particular pathway (or string), 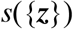. In this study, we explored the minimum free energy path of the transport pathway with the string method with the swarms of trajectories (SMwST)(Pan et al., 2008) in a two-dimensional collective variable space, (ΔD, Z_Pi_). To achieve the conformational sampling along the refined transition path by the bias exchange umbrella sampling (BEUS)(Moradi et al., 2015; Moradi & Tajkhorshid, 2014; Sugita et al., 2000) (simulation sets 6 and 12), the so-called path collective variables (pathCV)(Branduardi et al., 2007) were used:

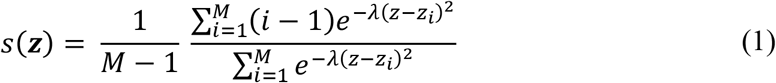

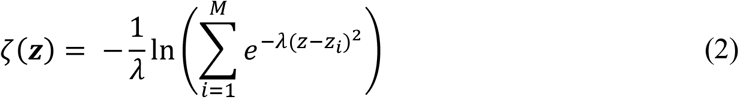

where λ is the average RMSD between images, *M* is the number of images, 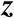 is a vector for the current values of the collective variables, and 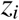 is the collective variables vector for the *i*-th image. In this paper, 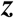 was represented as a two-dimensional vector (ΔD, Z_Pi_).

### Sampling protocol

At first, to generate the IF_b_^cent^ and OF_b_^cent^ states from the Occ state crystal structure, we performed a non-equilibrium pulling simulation (or steered MD simulation(Izrailev, Stepaniants, Balsera, Oono, & Schulten, 1997)) using the collective variable ΔD (simulation sets 1 and 2). These simulations were performed by moving the center harmonic restraint with a force constant of 10 kcal mol^−1^ Å^−2^ and using the ΔD ranges from 0 to −10 Å and 0 to 10 Å, to generate IF_b_^cent^ and OF_b_^cent^, respectively. The final structures from simulation sets 1 and 2 were used as the initial structures of the simulation sets 3 and 4, respectively, in which we induced the conformational transitions to IF_b_^intra^ and OF_b_^extra^ using the collective variable Z_Pi_ within the ranges from 0 to −15 Å and 0 to 15 Å, respectively.

As the IF_b_^intra^ ↔ OF_b_^extra^ transition was expected to undergo complicated conformational changes, including large scale substrate translocation and a significant local conformational change of the protein, we first refined the IF_b_^intra^ ↔ OF_b_^extra^ transition with SMwST, prior to reconstructing the entire conformational transition, IF_a_ ↔ Occ ↔ OF_a_. The final frames of these simulation sets were used to generate the initial string of the IF_b_^intra^ ↔ OF_b_^extra^ transition. We defined the initial IF_b_^intra^ ↔ OF_b_^extra^ transition string with 40 images: 10 frames from simulation set 3 (Z_Pi_ = 0, −1.5, −3.0, ···, −15 Å), 10 frames from simulation set 4 (Z_Pi_ = 0, 1.5, 3.0, ···, 15 Å) and 20 frames from simulation set 2 (ΔD = −10, −9, ···, 10 Å). This initial string was refined with SMwST in the two-dimensional (ΔD, Z_Pi_) collective variable space (simulation set 5). The selected images were minimized with 10,000 steps, followed by 1 ns of equilibration in the NPT ensemble with ΔD and Z_Pi_ restraints to the initial values. Twenty swarms of 0.1 ps unrestrained simulation were followed by 20 ps of restrained simulation, and the force constants for both ΔD and Z_Pi_ were 1.0 kcal mol^−1^ Å^−2^. The string was converged after 234 iterations. The last 20 strings were averaged and used as the IF_b_^intra^ ↔ OF_b_^extra^ transition path in simulation set 6. To perform the free energy calculation with BEUS along the refined transition string, we utilized the path collective variables(Branduardi et al., 2007) (pathCV(s) and pathCV(ζ) defined by equations (1) and (2) with the lambda value of 1.0. The 20 windows for the collective variable s = 0.025, 0.075, ··· 0.975 (increased by 0.05) were used for BEUS with a harmonic force constant of 200 kcal mol^−1^ Å^−2^. The collective variable ζ was restrained to 0.0 with a harmonic force constant of 200 kcal mol^−1^ Å^−2^. The replica exchange was attempted between neighboring replicas with an interval of 2.0 ps for each unique pair. The atomic coordinates were sampled every 2.0 ps.

Next, to induce the conformational transition to the *apo* structure, the final frames of simulation sets 1 and 2 were further employed in the non-equilibrium pulling simulation, using the collective variable Z_Pi_ (simulation sets 7 and 9) with the ranges from 0 to −45 Å and from 0 to 45 Å, respectively. BEUS with 16 windows along Z_Pi_ for the transition from IF_b_^cent^ to IF_a_ was performed with window centers (and force constants) of −2.0, −4.5, −6.5, −8.0, −9.0, −10.5, −12.0, −14.0, −15.5, −18.0, −20.5, −22.0, −24.0, −28.0, −33.0 and −40.0 Å (0.2, 0.5, 1.0, 1.0, 1.0, 0.5, 0.5, 0.5, 0.5, 0.5, 0.5, 0.5, 0.1, 0. 1, 0.05 and 0.01 kcal mol^−1^ Å^−2^) (simulation set 8). BEUS with 16 windows along Z_Pi_ for the transition from OF_b_^cent^ to OF_a_ was also performed with window centers and force constants of 0, 2.5, 4.0, 6.0, 8.5, 10.5, 12.5, 14.0, 16.0, 18.0, 20.0, 22.0, 25.0, 29.0, 33.0 and 40 Å (0.5, 1.0, 1.0, 0.5, 0.5, 0.5, 0.5, 0.5, 0.5, 0.5, 0.5, 0.5, 0.1, 0.1, 0.05 and 0.05 kcal mol^−1^ Å^−2^) (simulation set 10).

The structures and free energy information obtained from simulation sets 6, 8 and 10 were utilized for the PHSM(Moradi et al., 2015) to generate the initial string for the transition IF_a_ ↔ OF_a_. The PHSM was performed as described previously(Moradi et al., 2015). First, the initial string was generated by a non-parametric version of the lowest free energy pathway (LFEP)(Ensing, Laio, Parrinello, & Klein, 2005; Moradi et al., 2015). The following 37 metrics were used for the non-parametric LFEP and PHSM algorithm: (i) Z_Pi_, (ii) first 8 Cα-based principal components, (iii) D_in_ and D_out_, (iv) four atomic distances of K128-F192O, K128-V195O, K271-L347O and K271-V350O, (v) minimum distances (D_min_) between P_i_ and pore-lining polar residues: N124, K128, H201, K204, R266, K271, Y339, K362 and R363, (vi) D_min_ values of internal salt bridges E207-K204, E207-K362 and E207-R363, and (vii) χ_1_ angles for all 10 residues listed in (v) and (vi). A distance was defined in this 37-dimensional collective variable space of (i) to (vii), using a diagonal metric matrix with the values 1/Å^2^, 0.04/Å^2^, 1/Å^2^, 1/Å^2^, 0.01/Å^2^, 1/Å^2^ and 0.0025/(1°)^2^ for the elements in (i) to (vii), respectively. These weights were roughly chosen based on the relative variance of the collective variables. The initial string generated by the non-parametric LFEP was used for the PHSM algorithm with 400 image centers. The tube thickness ε = 0.5 was used in the PHSM algorithm, since it gives the smoothest pathway as compared with other results based on ε = 0.1, 0.2, … 1.0.

The string of the IF_a_ ↔ OF_a_ transition generated by the PHSM algorithm was further refined with SMwST (simulation set 11) in the two-dimensional collective variable space (ΔD, Z_Pi_). We selected 40 images from the string refined with the PHSM algorithm at even intervals. The selected images from PHSM were minimized and equilibrated in the same manner as the simulation set 5. In this analysis, 20 swarms of 5 ps unrestrained simulation were performed, followed by 10 ps of restrained simulation with force constants of 10.0 kcal/mol for each collective variable. The string was converged after 50 iterations. The final 10 strings were averaged, and used for the final BEUS sampling.

The final BEUS sampling along the refined IF_a_ ↔ OF_a_ transition string (simulation set 12) was performed using the pathCV(s) and pathCV(ζ) with λ = 0.1 (equations (1), (2)). The 40 images corresponding to the image points of the averaged string were selected and equilibrated for 0.1 ns with the collective variables (ΔD, Z_Pi_) restrained. In total, 40 windows, s = 0.015, 0.035, … 0.99 (increased by 0.025) for the pathCV(s) were used for BEUS, with a force constant of 500 kcal mol^−1^. The pathCV(ζ) is restrained to the region ζ < 1.0 with half-harmonic restraints with a force constant of 100 kcal mol^−1^. The exchange was attempted between the neighboring replicas with an interval of 10 ps for each unique pair. The sampling was performed for 50 ns, and the coordinates for all atoms were sampled every 2 ps, to obtain 1,000,000 structures (= 40 replicas × 50 ns / 2 ps) in total.

### Data analysis

Data analysis was performed using MDAnalysis(Michaud-Agrawal, Denning, Woolf, & Beckstein, 2011), mdtraj(McGibbon et al., 2015), and HOLE(Smart et al., 1996), along with in-house python and C++ codes. All molecular graphics were illustrated with CueMol2 (http://www.cuemol.org/). Plot graphics were generated with ggplot2(Wickham, 2009), seaborn(Waskom et al., 2016) and matplotlib(Hunter, 2007).

To calculate the potential of mean force (PMF) from the BEUS simulations along the path collective variable s in Figure 1f, the C implementation of the WHAM algorithm(Kumar et al., 1992) (Grossfield, Alan, “WHAM: the weighted histogram analysis method”, version 2.0.9 http://membrane.urmc.rochester.edu/content/wham) was used. To perform the *post-hoc* string method analysis, the weights for each structure *w*^*t*^ were required(Moradi et al., 2015). They were calculated by the following generalized WHAM equation(Bartels, 2000; Moradi et al., 2015):

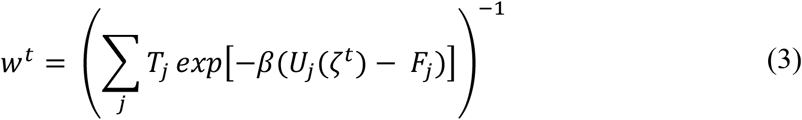

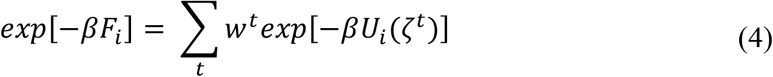

where *β* = 1/*k*_B_*T* is the inverse temperature, *T*_*j*_ is the number of samples collected for image *j*, *F*_*j*_ is the PMF for the *j*-th umbrella window, *ζ*^*t*^ is the collective variable at time *t*, and 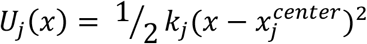 is the *j*-th umbrella potential used in the BEUS simulation. The projection of the PMF from one collective variable space to another one, ξ, was performed with the following equation(Moradi et al., 2015):

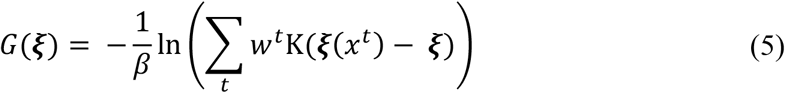

where K(.) is a kernel function. In this study, the RBF kernel was used.

The principal component analysis (PCA) was performed on the Cα atoms of GsGPT. The coordinates of the trajectory were superimposed on the averaged structure of simulation set 12. The covariance matrix of the atomic fluctuations was calculated, and its eigenvalues and eigenvectors were determined. The calculation of the number of hydrogen bonds in Figure 3 was performed by the following equation:

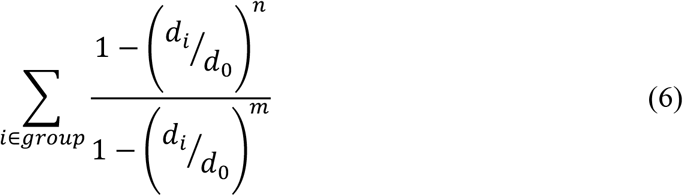

where **d**_*i*_ is the distance between two hydrogen-bonding atoms, *d*_0_ is a cutoff distance, and *n* and *m* are integers that determine the behavior of the switching function. The sum is taken over all of the hydrogen bonding atom pairs. In this study, *d*_0_ = 3.3 Å, *n* = 6 and *m* = 8 were used.

### Transport assay

The liposome assays were performed as previously described(Lee et al., 2017). In brief, the liposomes reconstituted with yeast membranes expressing WT or mutant GsGPT were preloaded with 30 mM NaH_2_PO_4_, and were mixed with an equal volume of extra-liposomal solution containing 1 mM [^32^P]-NaH_2_PO_4_ (0.1 mCi ml^−1^). The reaction was terminated by passing the liposomes through AG-1 X8 resin. The total amounts of incorporated P_i_ were measured at 30 min. The control experiment was performed with membranes from yeast cells harboring the empty vector. The mean count value of the control experiment was subtracted from each count value for each experiment set, and the values shown in the figures were normalized with the mean value of the WT. The expression of each mutant was verified by a western-blot analysis, using an anti-His-tag polyclonal antibody (code PM032; MBL).

## Acknowledgements

We thank R. Taniguchi and H. Miyauchi (University of Tokyo, Japan) for comments on the manuscript, and T. Nakane (University of Tokyo, Japan) for technical assistance. The *post-hoc* string method program was kindly provided by Dr. Mahmoud Moradi (University of Arkansas). Computations of MD simulations were partially performed on HOKUSAI GreatWave at the RIKEN Advanced Center for Computing and Communication, the mini-K super computer system at the SACLA facility, and the NIG supercomputer at ROIS National Institute of Genetics. This work was supported by a Grant-in-Aid for Specially Promoted Research (16H06294) and a Grant-in-Aid for Scientific Research (B) (25291011) from the Japan Society for the Promotion of Science (JSPS) to O.N. and R.I., respectively.

## Author contributions

M.T. designed the research, performed the molecular dynamics simulation, and analyzed the data. Y.L. performed the liposome transport assay. M.T., R.I. and O.N. wrote the manuscript. R.I. and O.N. directed and supervised all of the research.

---

**Figure 1—figure supplement 1.**
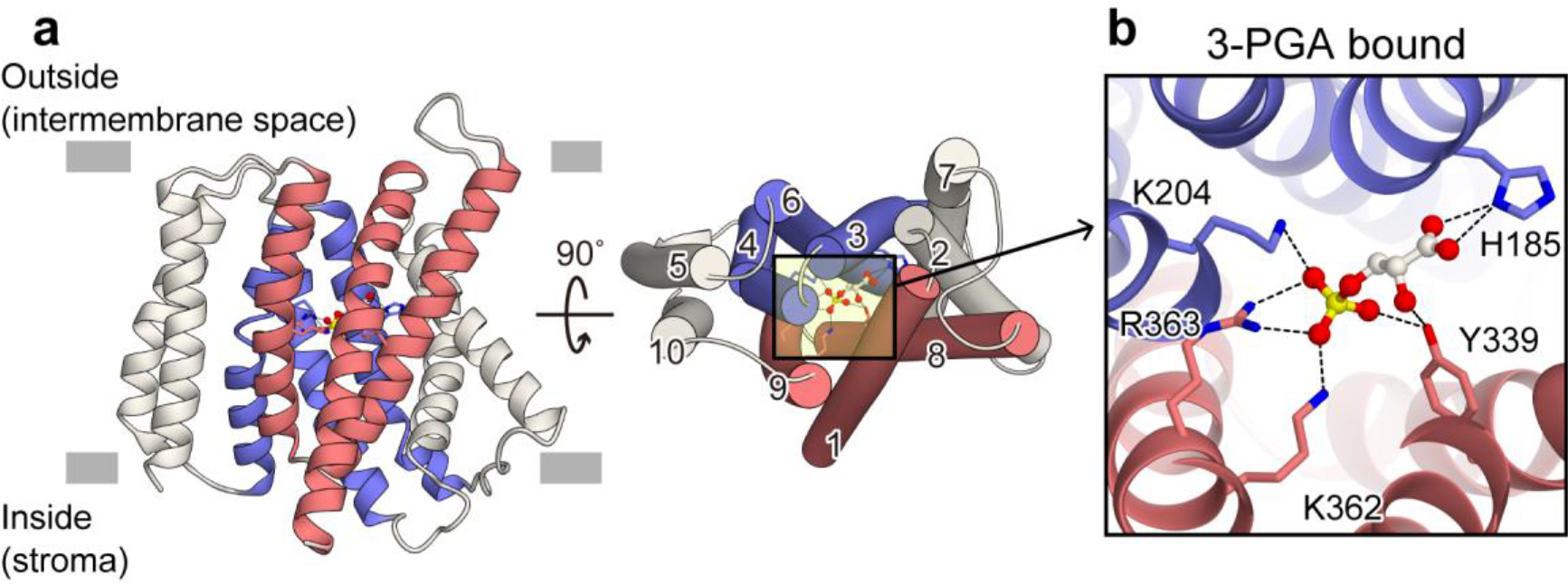
Crystal structure of GsGPT with 3-PGA. (a) The crystal structure of GsGPT in the 3-phosphoglyceric acid (3-PGA)-bound state (PDB ID: 5Y79), viewed from the plane of the membrane (left) and from the intermembrane space (right). The numbers in the right panel represent the numbering of the TM helices. TM1, 8 and 9 are colored red, TM3, 4 and 6 are colored blue, and the others are colored white. (b) Close-up view of the central binding site in the 3-PGA-bound state. Key residues involved in substrate binding are shown in stick models. Dotted lines represent polar interactions.

**Figure 1—figure supplement 2.**
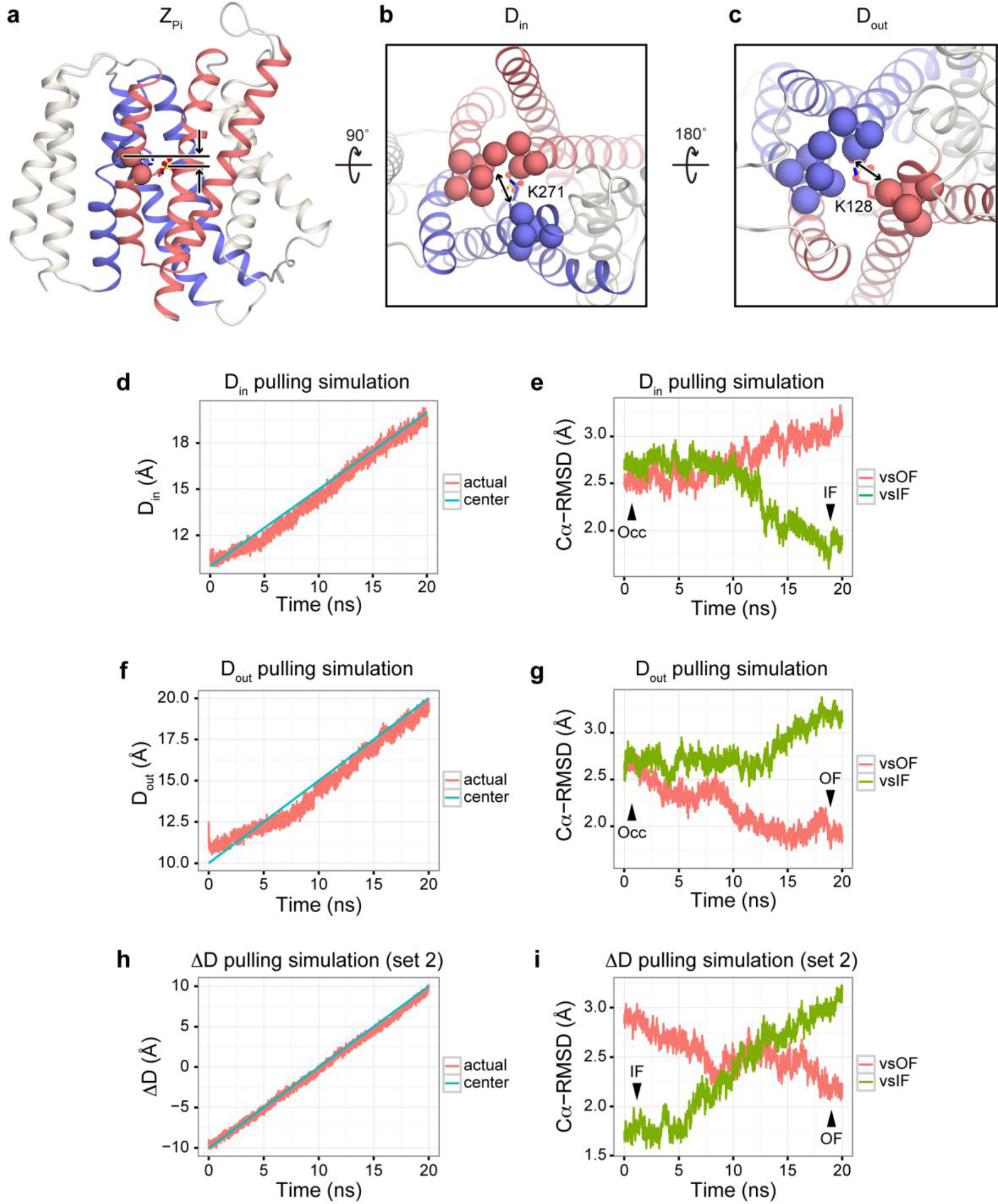
The definitions of collective variables and the steered MD simulations. (a-c) Definitions of collective variables. (a) Definition of Z_Pi_. P_i_ is shown with a ball and stick model, and the Cα atoms of Lys204, Lys362 and Arg363 used for the
reference points are shown with spheres. (b, c) Definitions of D_in_ and D_out_. The Cα atoms used for the calculation are shown as spheres. (d, f, h) Time courses of the collective variables, (d) D_in_, (f) D_out_ and (h) ΔD, for the non-equilibrium pulling (steered MD) simulation of each variable. The blue line represents the center of the harmonic potential, and the red line represents the actual value of the collective variable. (e, g, i) Time courses of the root-mean-square deviation (RMSD) values of all Cα atoms relative to the OF (red) and IF (green) conformations. IF, Occ and OF represent the initial, occluded and final conformations of the protein, respectively.

**Figure 2—figure supplement 1.**
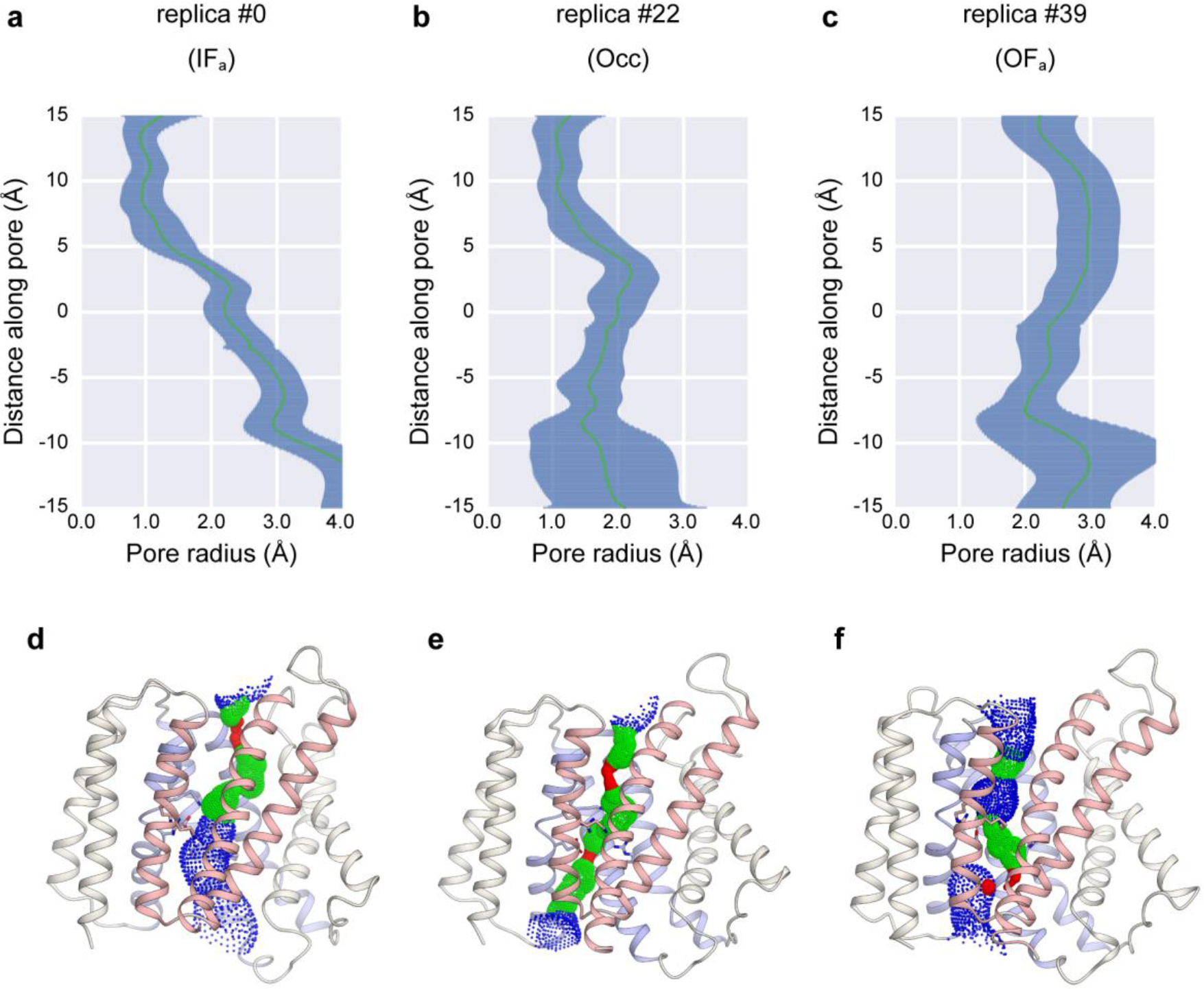
Representative results for pore radius calculation by HOLE. (a-c) Minimum radius along the path through the central binding site for (a) replica#0 (IF), (b) replica#22 (Occ) and (c) replica#39 (OF). The green line represents the mean value and the blue line represents the mean value and standard deviation for each distance along the pore. (d-f) Representative structures of the (d) IF, (e) Occ and (f) OF states. The red region represents *r* < 0.6 Å, the green region represents 0.6 < *r* < 1.2 Å and the blue region represents *r* > 1.2 Å.

**Figure 2—figure supplement 2.**
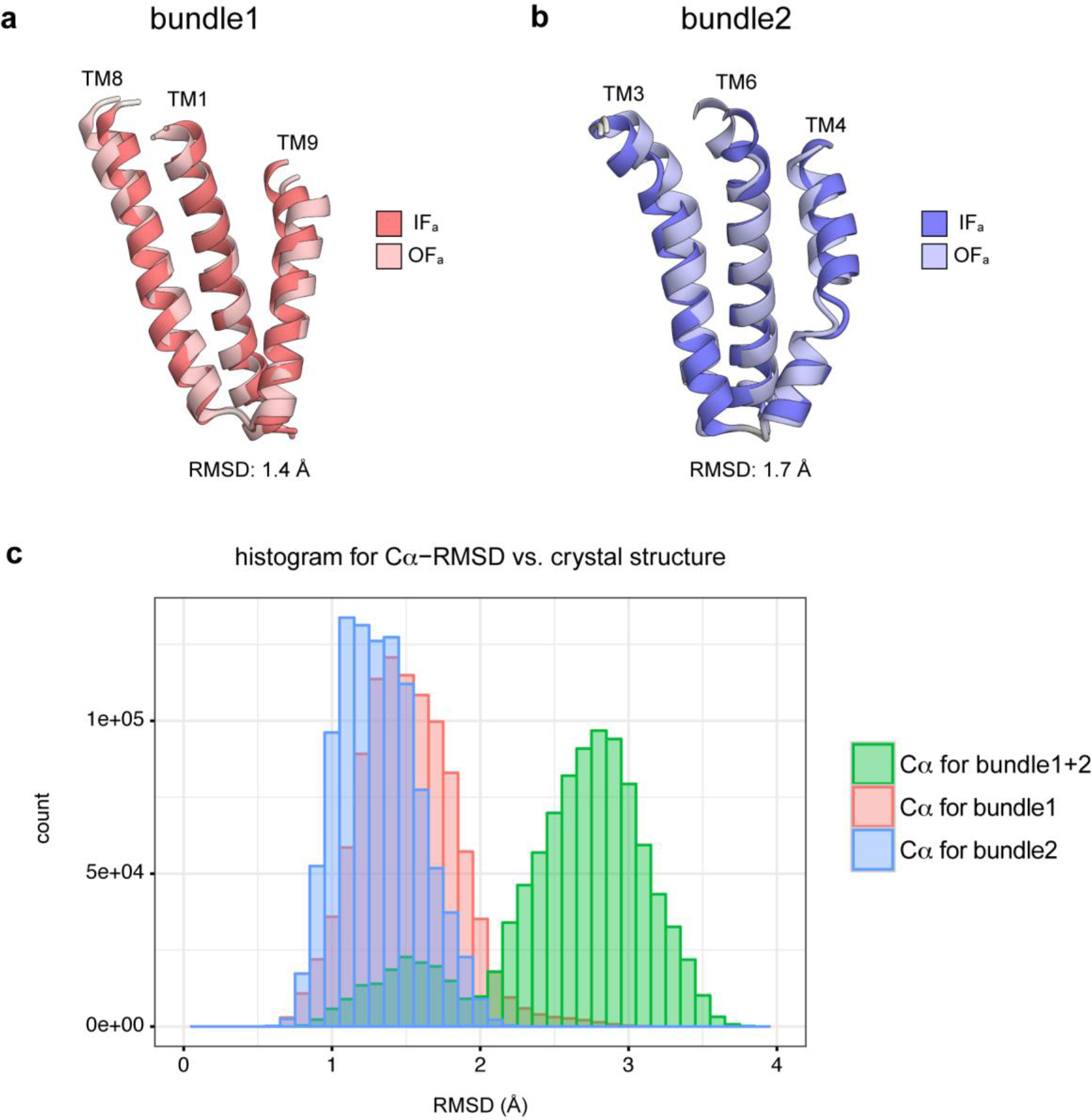
Intra-domain movement of the core domain. (a, b) Representative conformations of (a) bundle1 and (b) bundle2 from the IF and OF conformations. The root-mean-square deviation (RMSD) values of the Cα atoms for each structure are shown below. (c) The histogram of RMSD values of Cα atoms in bundles1+2 (green), bundle1 only (red) and bundle2 only (blue) values relative to the crystal structure. The RMSD values were calculated for the α-helix region of each bundle.

**Figure 2—figure supplement 3.**
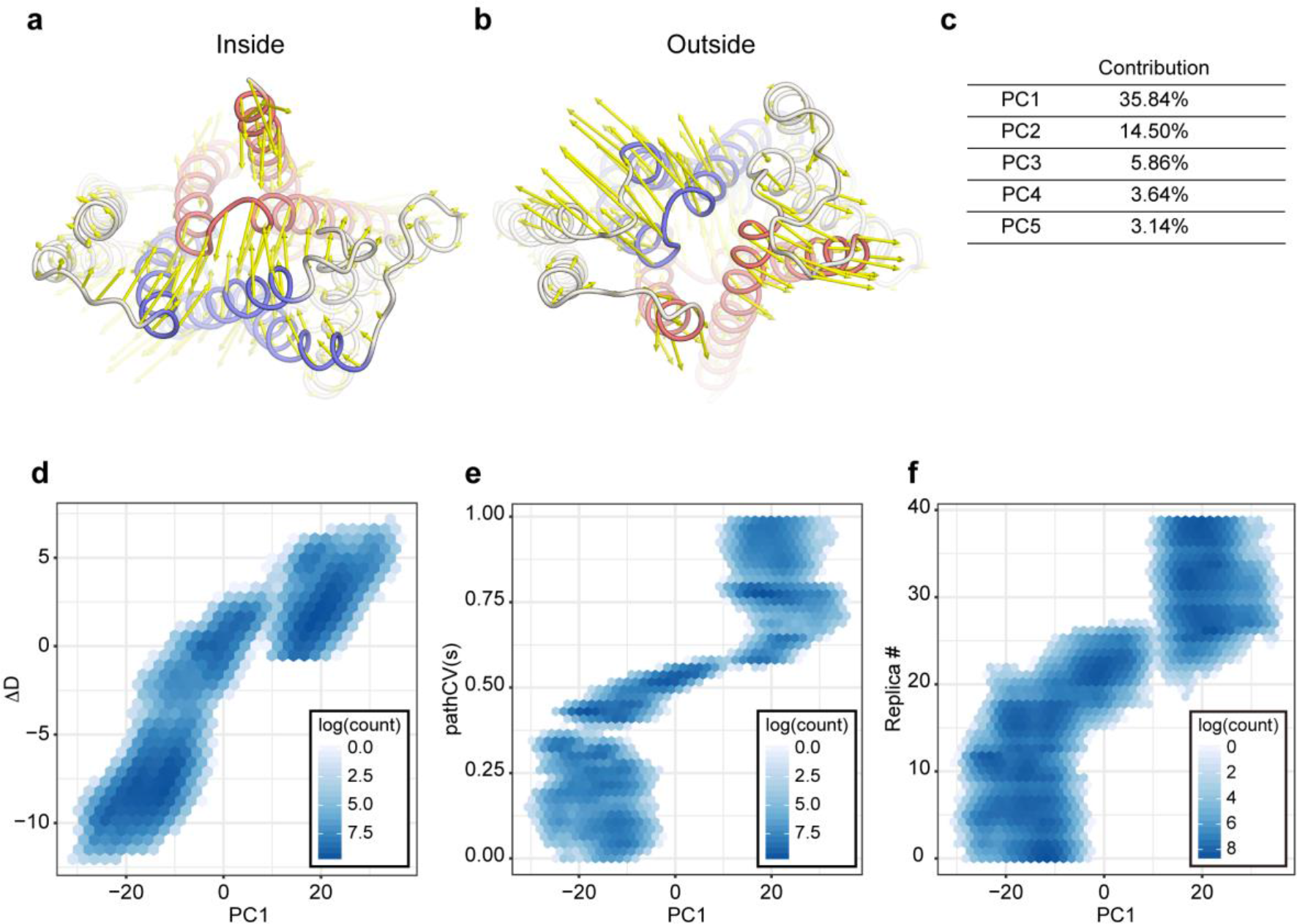
The visualization of PCA results and correlations with other variables. (a, b) Vector representation of the eigenvector of the first principal component (PC1) of GsGPT movement, calculated by principal component analysis (PCA). (c) The contributions of PC1-5 for the overall deviations. (d-f) Correlations of the PC1 *vs*. (d) ΔD, (e) the number of replicas and (f) the pathCV(s) shown in a 2D histogram. The count value is shown in a log scale.

**Figure 3—figure supplement 1.**
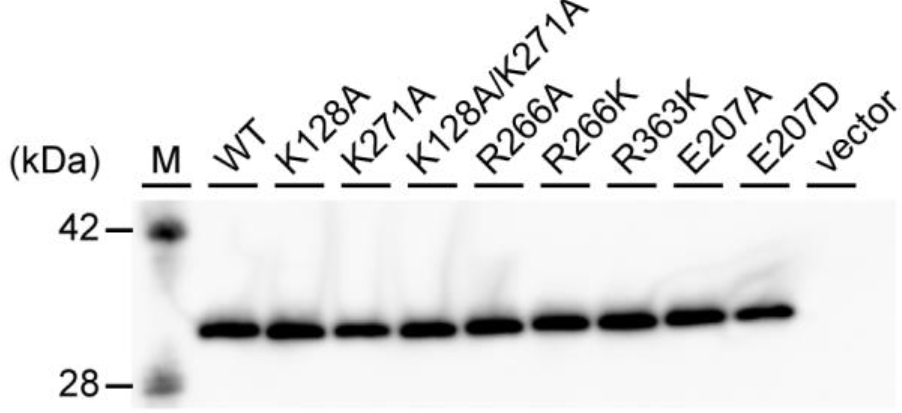
Membrane expression of GsGPT mutants. Western blotting analysis for wild type (WT) and each GsGPT mutant, confirming the comparable expression levels. “M” represents marker lane.

**Figure 4—figure supplement 1.**
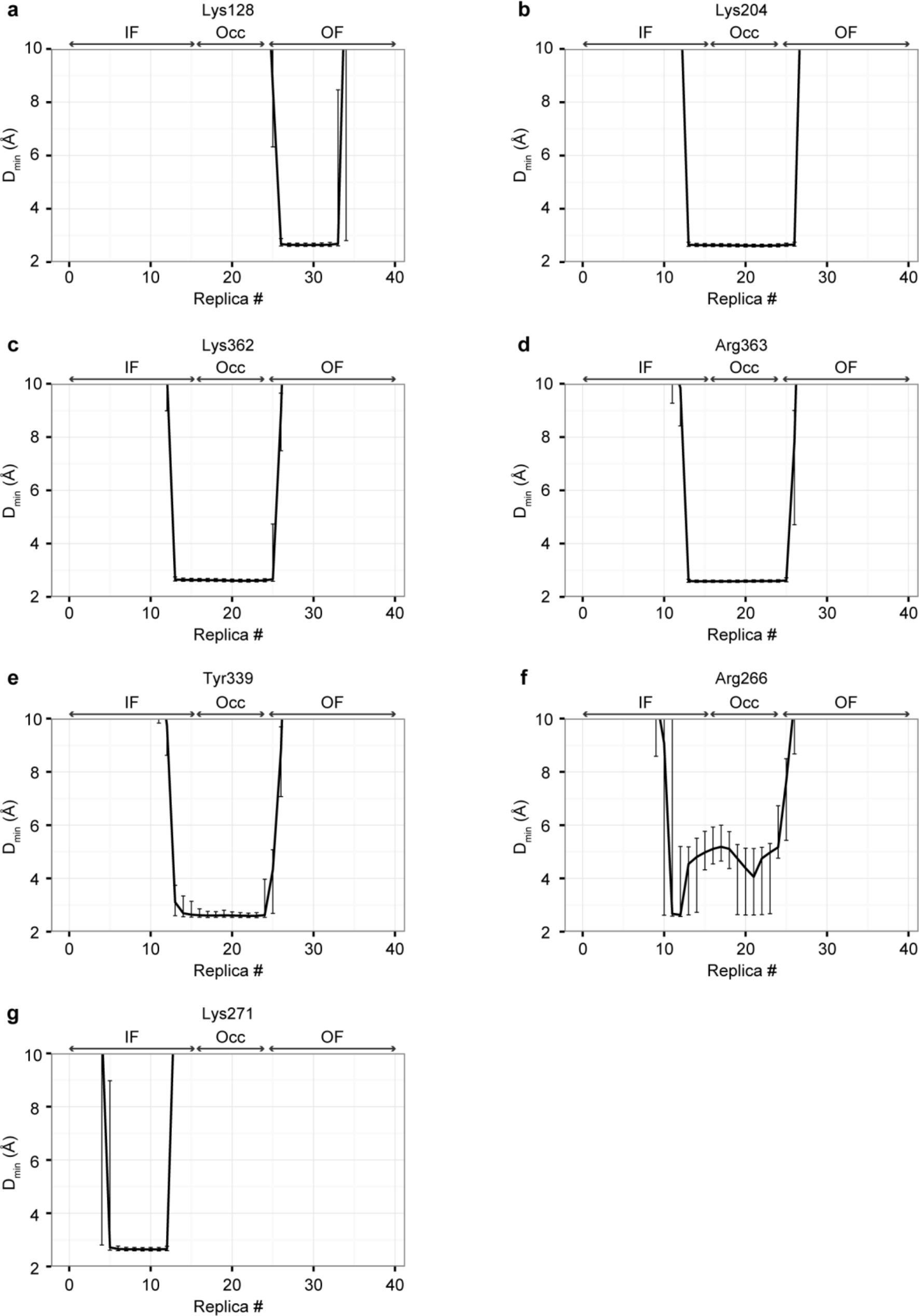
Minimum distances between P_i_ and the P_i_ interacting residues. The minimum distances between P_i_ and the P_i_ interacting residues, (a) Lys128, (b) Lys204, (c) Lys362, (d) Arg363, (e) Tyr339, (f) Arg266 and (g) Lys271. The median value within each replica is plotted, and the error bars represent the interquartile range (IQR).

**Figure 4—figure supplement 2.**
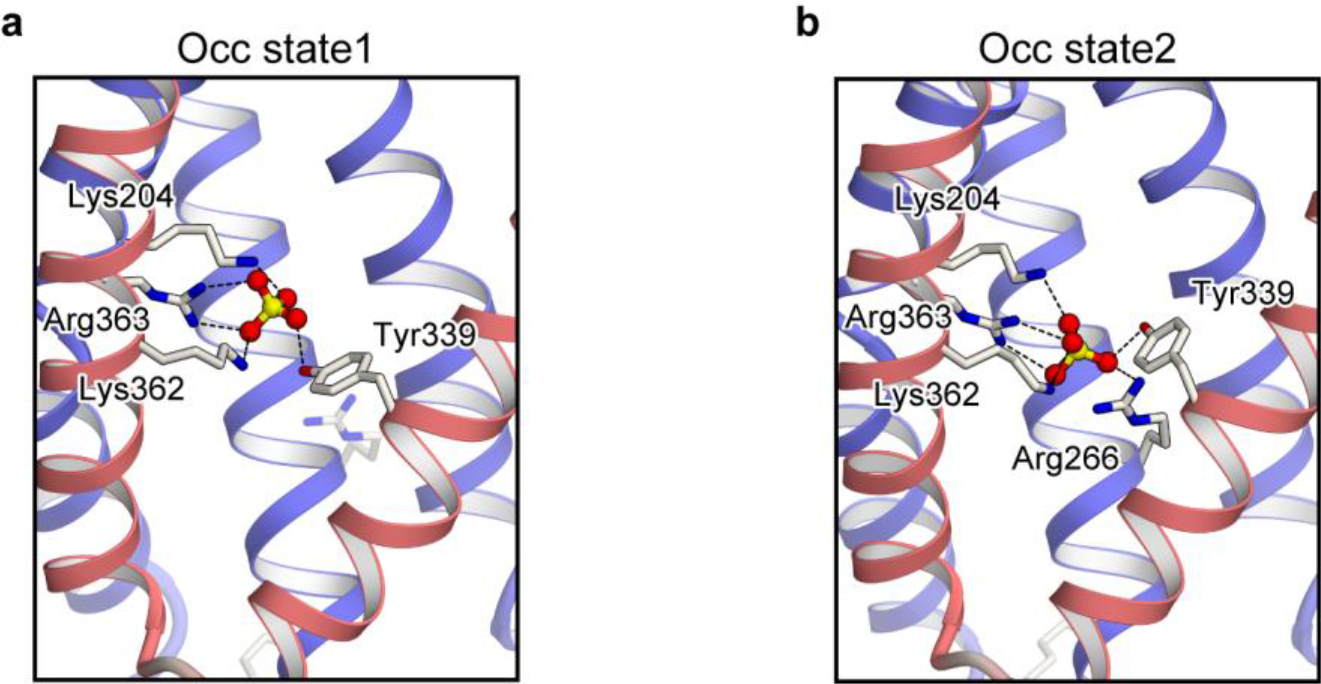
Degenerate states in the Occ conformation. (a, b) Close-up views of the substrate binding site for two conformations corresponding to the free energy basins shown in Figure 4d. TM1 is not shown.

**Figure 5—figure supplement 1.**
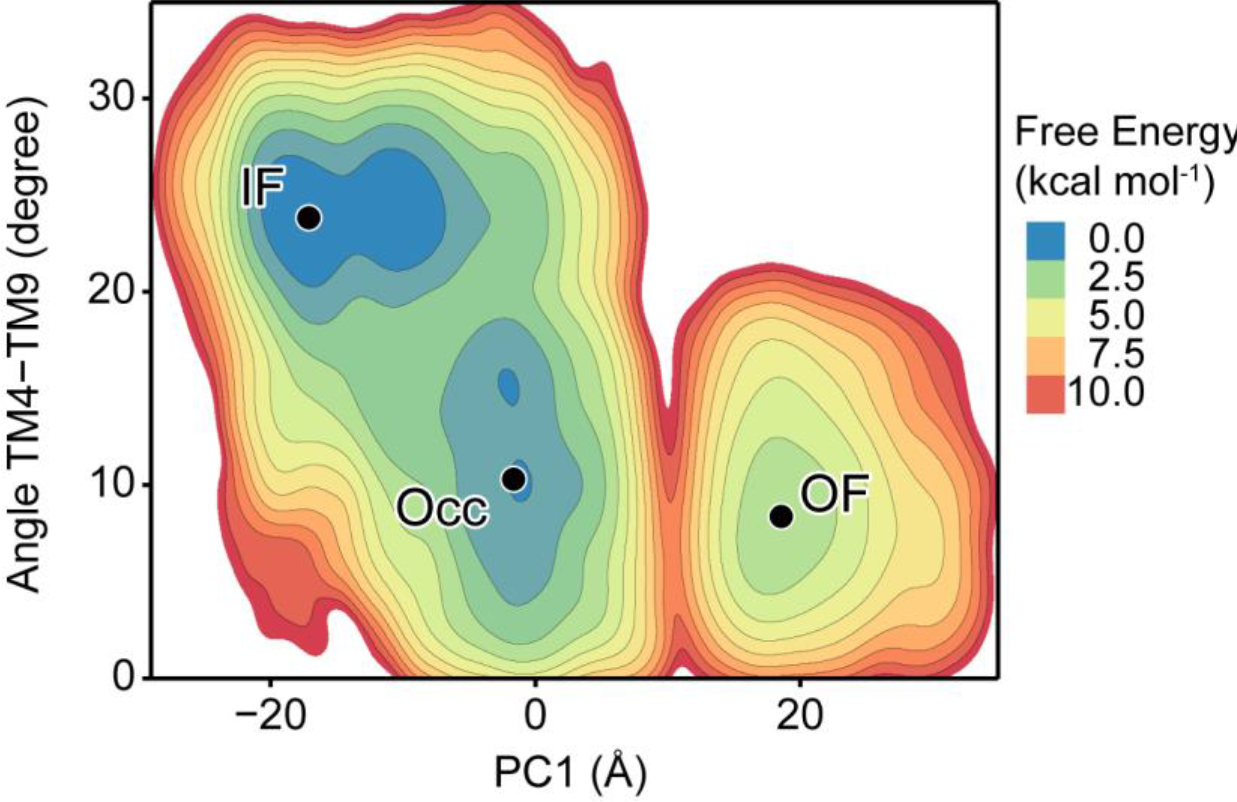
Free energy landscape for inter-helical angle between TM4 and TM9. Free energy landscape in terms of (PC1, TM4-TM9 angle) space. PC1 represents the first principal component of all Cα atoms, and the inter-helical angle between TM4 and TM9 was calculated using the roll axis of the helix (obtained from the principal axis component analysis for the Cα atoms).

## Rich media files

### Video1

IF-to-OF conformational transition of GsGPT, viewed from the plane of the membrane. The trajectory was reconstructed by the *post-hoc* string method (PHSM) from the final bias-exchange umbrella sampling (BEUS) simulation (simulation set 12). The P_i_ molecule is shown with a ball-and-stick model, and the major substrate binding basic residues and Glu207 are highlighted with stick models. The coloring scheme for the TM helices is the same as in Figure 1.

### Source Data

Figure 3—Source Data1 | List of the raw count values of transport assay

